# Alzheimer’s Disease Risk Variants Interact with Amyloid-beta to Modulate Monocyte Function

**DOI:** 10.1101/2025.11.02.686117

**Authors:** Zena K. Chatila, Elizabeth M. Bradshaw

## Abstract

While genetics implicate a central role for dysregulated innate immunity in Alzheimer’s disease (AD), the contributions of peripheral myeloid cells, such as monocytes, have been largely overlooked in favor of microglia. Here, we investigate whether AD associated loci, specifically *rs3865444* in the *CD33* locus and *rs1057233* in the *SPI1* locus, converge on shared functional pathways in monocytes in the context of amyloid-beta peptide 1-42 (Aβ1-42) as an immune stimulus. To do so, we isolated monocytes from peripheral blood mononuclear cells (PBMCs) from healthy individuals and exposed them to aggregated Aβ1-42. In this study, we identify functional convergence of the *CD33* and *SPI1* AD risk variants in the context of aggregated Aβ, both resulting in reduced phagocytosis and loss of surface TREM2 expression, demonstrating an interaction between genetics and environment to reduce myeloid cell fitness. These findings highlight that peripheral monocytes, like brain-resident microglia, are genetically and functionally linked to AD risk, underscoring their importance as accessible immune cells that contribute to disease susceptibility and progression.

## Introduction

Alzheimer’s disease (AD) is an age-related neurodegenerative disease characterized by progressive cognitive decline as well as pre-symptomatic accumulation of amyloid pathology with older age. With the discovery and validation of AD susceptibility loci, several risk factors have been identified that are solely or highly expressed in innate immune myeloid cells^1^. Such discoveries have highlighted risk genes for the disease including *CD33, TREM2*, and *SPI1*, and implicate a central role for myeloid dysfunction in AD. Importantly, many of these genetically associated proteins interact with each other and function in shared pathways^1^. However, the functional significance of these risk variants on peripheral myeloid cell activity in AD remains unclear.

While the discovery of AD genetic risk variants enriched in myeloid gene loci has led to a dedicated effort to understand microglial involvement in AD pathogenesis, myeloid cells in the periphery have been largely overlooked. Importantly, genetic data implicating myeloid cell involvement does not discriminate between those that are resident in the CNS parenchyma, such as microglia and perivascular macrophages, and those that are found in the periphery, such as monocytes and macrophages^2^. Not only can peripheral myeloid cells infiltrate the CNS, but they can also alter immune dynamics in the periphery itself and influence the adaptive immune system. Indeed, peripheral myeloid cells are even thought to impact clearance of amyloid beta (Aβ) in the CNS, both by trafficking to the CNS and participating in Aβ clearance locally, as well as through phagocytosing Aβ in the periphery, which has in turn been found to alter CNS Aβ clearance^3,4^. Increasing evidence has also implicated a role for monocyte homing to the brain and compensating for microglial dysfunction in AD^5^. While both cell types are involved in amyloid clearance, monocytes are thought to have significantly higher phagocytic capacity than microglia^6^. Thus, understanding the functional impact of AD genetic risk factors on monocyte function is valuable for identifying pathways through which genetic variants modulate disease risk and uncovering novel therapeutic approaches for this disease.

The present work aims to clarify the phenotypic consequences of the interaction between AD-associated loci enriched in myeloid cells and the response to Aβ in monocytes. CD33 is an inhibitory transmembrane receptor expressed on myeloid cells. The risk variant *CD33 rs3865444*^*C*^ associated with AD results in increased CD33 surface expression, reduced phagocytosis, and an increased amyloid burden in the brain^7^. The *CD33 rs3865444*^*C*^ AD risk variant not only increases CD33 surface expression but also increases TREM2 surface expression^8^. TREM2 is an excitatory surface receptor expressed exclusively on myeloid cells, which signals through the associated adaptor protein DAP12 (encoded by *TYROBP*), and has been associated with AD through whole-exome sequencing. Several studies have demonstrated that AD-associated genetic variants in TREM2 result in impaired signaling, suggesting that a partial loss of TREM2 function may be a major contributor to AD pathogenesis^1,9^. CD33 regulation of TREM2 surface expression is specifically mediated by membrane-bound CD33; individuals with the *CD33* AD risk variant have increased surface TREM2 expression, an effect which was abrogated with a CD33 neutralizing antibody which downregulates surface CD33^8^. TREM2 has also been found to function downstream of CD33 in mouse studies utilizing the 5xFAD model^10^.

*CD33* and *TREM2* gene expression are both regulated by the transcription factor PU.1 (encoded by *SPI1*), which has also been genetically associated with AD through GWAS^11^. PU.1 functions as a master regulator transcription factor in myeloid cells^11^. The protective *rs1057233*^*G*^ variant associated with reduced AD risk results in reduced PU.1 expression. Notably, PU.1 regulates the expression of several AD-associated genes, including *CD33* and *TREM2*, as well as *TYROBP*^*1,11-13*^. Specifically, PU.1 expression is necessary for CD33 expression in myeloid cells, and silencing of *SPI1* with siRNA causes reduced TREM2 expression^12,13^. Like *rs3865444* in the *CD33* locus, *rs1057233* in the *SPI1* locus has been connected to phagocytic capacity of innate immune cells. However, knockout of *SPI1* in microglia surprisingly resulted in decreased phagocytosis in BV2 cells, a model of murine microglia^14^.

The goal of this work is to determine whether these functionally interconnected AD-associated loci and proteins converge to alter monocyte function in the context of an AD-associated stressor, aggregated Aβ. To do so, we isolated monocytes from PBMCs from healthy individuals and exposed them to aggregated Aβ1-42. We identify convergence of AD loci on shared functional outcomes in monocytes in the context of amyloid as an immune stimulus, including reduced phagocytosis and loss of surface TREM2 expression. These findings further support a role for TREM2 loss of function in myeloid dysfunction in AD. While immune-targeting therapies in development are designed to cross the blood brain barrier to target microglial cells within the brain, future work may clarify whether modulating TREM2, CD33, or PU.1 signaling in the periphery alone may be beneficial in AD^15^.

## Results

### Aggregated Aβ1-42 Impairs Phagocytosis in Monocytes Carrying *CD33* and *SPI1* AD Risk Variants

To examine the interaction between AD genetic risk variants and environmental stimuli associated with disease pathology in the periphery, we isolated monocytes from individuals genotyped for *SPI1 rs1057233* and *CD33 rs3865444* AD risk variants (Supplemental Table 1) and exposed them to aggregated Aβ1-42 *in vitro* for 24 hours to provide a disease-associated stressor. As CD33 and PU.1 AD variants have both been associated with phagocytosis, we first examined whether phagocytic function of monocytes is altered following exposure to aggregated Aβ1-42^7,11^. We measured phagocytosis of fluorescently labeled Aβ1-42 with flow cytometry. In aggregate (across all genotypes combined), exposure to aggregated Aβ1-42 had no significant impact on phagocytosis (**Fig. 1a, Extended Data Fig. 1a**). However, when we examined phagocytosis in a genotype-specific manner, we found that aggregated Aβ1-42 treatment resulted in impaired phagocytosis only in monocytes from individuals that were homozygous for either the *CD33* or *SPI1* AD risk variants, revealing a functional impairment dependent on the interaction between genetics and environment (**Fig. 1b, 1c, Extended Data Fig. 2a**). No differences in phagocytosis were observed in association with sex or age (Extended Data Fig. 3, Extended Data Fig. 4).

**Figure 1.**
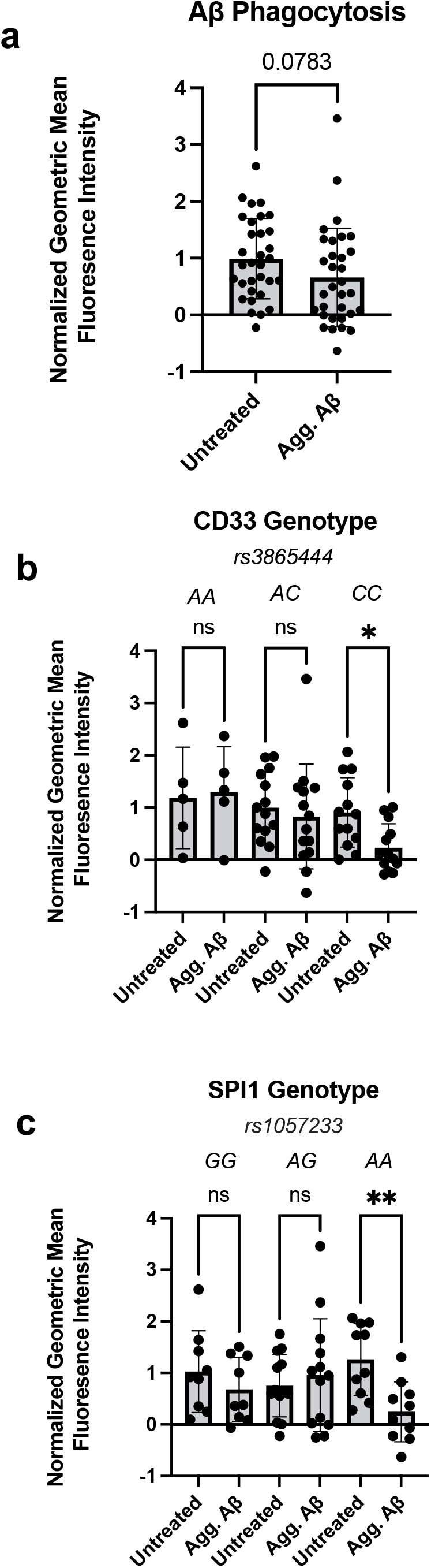
Aggregated Aβ1-42 Treatment Reduces Phagocytosis of Amyloid Beta in Monocytes Carrying *CD33* and *SPI1* Alzheimer’s Disease Risk Variants. **a**. Human monocyte phagocytosis of fluorescently labeled amyloid beta peptide 1-42 (Aβ1-42) following exposure to aggregated Aβ1-42 (Agg. Aβ) for 24 hours, measured with flow cytometry. Data are represented as geometric mean fluorescence intensity and normalized for batch correction using the GraphPad Prism normalize function. **b**. Monocyte phagocytosis of fluorescently labeled Aβ1-42 as in **a**, stratified by *CD33 rs3865444* genotype. *CD33 rs3865444*^*C*^ is associated with increased AD risk. **c**. Monocyte phagocytosis of fluorescently labeled Aβ1-42 as in **a**, stratified by *SPI1 rs1057233* genotype. *SPI1 rs1057233*^*A*^ is associated with increased AD risk. **a-c**: Data is presented as the mean ± SEM, and each dot represents an individual donor. **a**: paired Student’s t-test, **b-c:** two-way ANOVA with Sidak’s multiple comparison test, *p<0.05, **p<0.01.

Importantly, monocytes can uptake Aβ through phagocytosis-independent mechanisms. We performed these experiments with the addition of Cytochalasin D, which inhibits phagocytosis, and found reduced Aβ1-42 internalization **(Extended Data Fig. 1a**). These findings confirm that the uptake we observed occurs through phagocytosis.

### Aggregated Aβ1-42 Reduces Monocyte TREM2 Surface Expression in *CD33* and *SPI1* Risk Variants

TREM2 surface expression is functionally implicated in phagocytosis^16-18^. CD33 regulates TREM2 surface expression^8,10^, and PU.1 regulates *TREM2* gene expression^12^, suggesting alterations in TREM2 may be one shared factor modulating the altered phagocytosis we observed with these risk variants. Thus, we next investigated whether the interaction between CD33 or PU.1 and TREM2 surface expression is altered in the context of aggregated Aβ1-42. We exposed monocytes isolated from PBMCs of healthy individuals genotyped for *CD33* and *SPI1* AD risk variants to aggregated Aβ1-42 *in vitro* for 24 hours and measured surface TREM2 expression with flow cytometry.

In aggregate, across all genotypes combined, aggregated Aβ1-42 treatment resulted in reduced TREM2 surface protein expression, regardless of the genotype of the individual (**Fig. 2a, Extended Data Fig. 1b**). However, when we analyzed these findings in a genotype-specific manner, we determined this effect was driven by those who were homozygous risk for either the *CD33* or *SPI1* AD variants, as only these stratified groups demonstrated significantly reduced TREM2 surface expression (**Fig. 2b, 2c**). This specific reduction in surface TREM2 again demonstrates a genotype-dependent response to aggregated Aβ1-42. Importantly, the *CD33* genotype has been shown to modulate TREM2, such that the *rs3865444*^C^ risk variant results in increased TREM2 surface expression^8^. We observed this same effect at baseline (**Extended Data Fig. 2b**), and found that, following aggregated Aβ1-42 treatment, monocytes homozygous for *rs3865444*^*CC*^ no longer had increased surface TREM2 expression compared to *rs3865444*^*CA*^ and *rs3865444*^*AA*^ genotypes.

**Figure 2.**
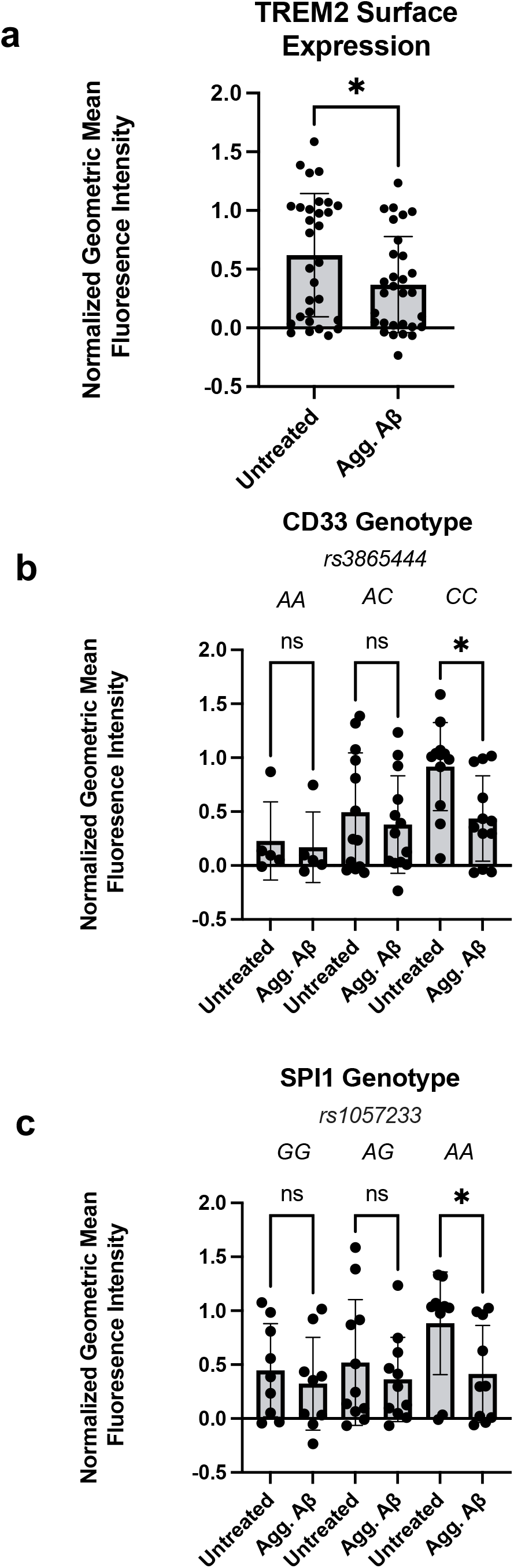
Aggregated Aβ1-42 Treatment Reduces TREM2 Surface Expression in Monocytes Carrying *CD33* and *SPI1* Alzheimer’s Disease Risk Variants. **a**. Human monocyte surface expression of TREM2 following exposure to aggregated amyloid beta peptide 1-42 (Agg. Aβ) for 24 hours, measured with flow cytometry. Data are represented as geometric mean fluorescence intensity and normalized for batch correction using the GraphPad Prism normalize function. **b**. Monocyte TREM2 surface expression as in **a**, stratified by *CD33 rs3865444* genotype. *rs3865444*^*C*^ is associated with increased AD risk. **c**. Monocyte TREM2 surface expression as in **a**, stratified by *SPI1 rs1057233* genotype. *rs1057233*^*A*^ is associated with increased AD risk. Data are represented as geometric mean fluorescence intensity and normalized for batch correction with min-max scaling. **a-c**: Data is presented as the mean ± SEM, and each dot represents an individual donor. **a**: paired Student’s t-test, **b-c:** two-way ANOVA with Sidak’s multiple comparison test, *p<0.05, **p<0.01.

We next investigated whether TREM2 surface expression is mediated by age or sex. No differences in TREM2 were observed in association with age (Extended Data Fig. 4b). Males had decreased surface TREM2 following treatment with aggregated Aβ1-42, while females had no difference in TREM2 expression with Aβ1-42 exposure (**Extended Data Fig. 3b**).

### Aggregated Aβ1-42 Influences *TREM2* and *TYROBP* Transcription in a Genotype-Independent Manner

As a key innate immune transcription factor, PU.1 regulates the transcription of both *CD33* and *TREM2*, as well as *TYROBP*^*1,11-13,19*^. To understand whether transcriptional alterations triggered by the response to aggregated Aβ1-42 may provide a convergent mechanism underlying the shared phenotype we observed between *CD33* and *SPI1* risk variants, we examined RNA expression of these genes in genotyped monocytes following aggregated Aβ1-42 stimulation.

We found that treatment with aggregated Aβ1-42 altered the transcription of *TREM2* and *TYROBP*. Specifically, both *TREM2* and *TYROBP* gene expression were decreased following aggregated Aβ1-42 treatment in monocytes across all genotypes combined (**Fig. 3a, 3b**). When stratified by *CD33* and *SPI1* genotype, this reduced *TREM2* and *TYROBP* gene expression did not follow a genotype-specific pattern (**Fig. 3c-f**). No genotype-specific differences were found in expression of either of these genes in untreated monocytes (**Extended Data Fig. 5**).The discordance between a reduction of *TREM2* gene expression in a genotype-independent manner and the specific risk genotype-dependent alterations in TREM2 surface protein levels following aggregated Aβ1-42 treatment are suggestive of differential post-translational regulation of TREM2 levels with the risk variants, such as intracellular sequestration, cleavage, or secretion of the protein.

**Figure 3.**
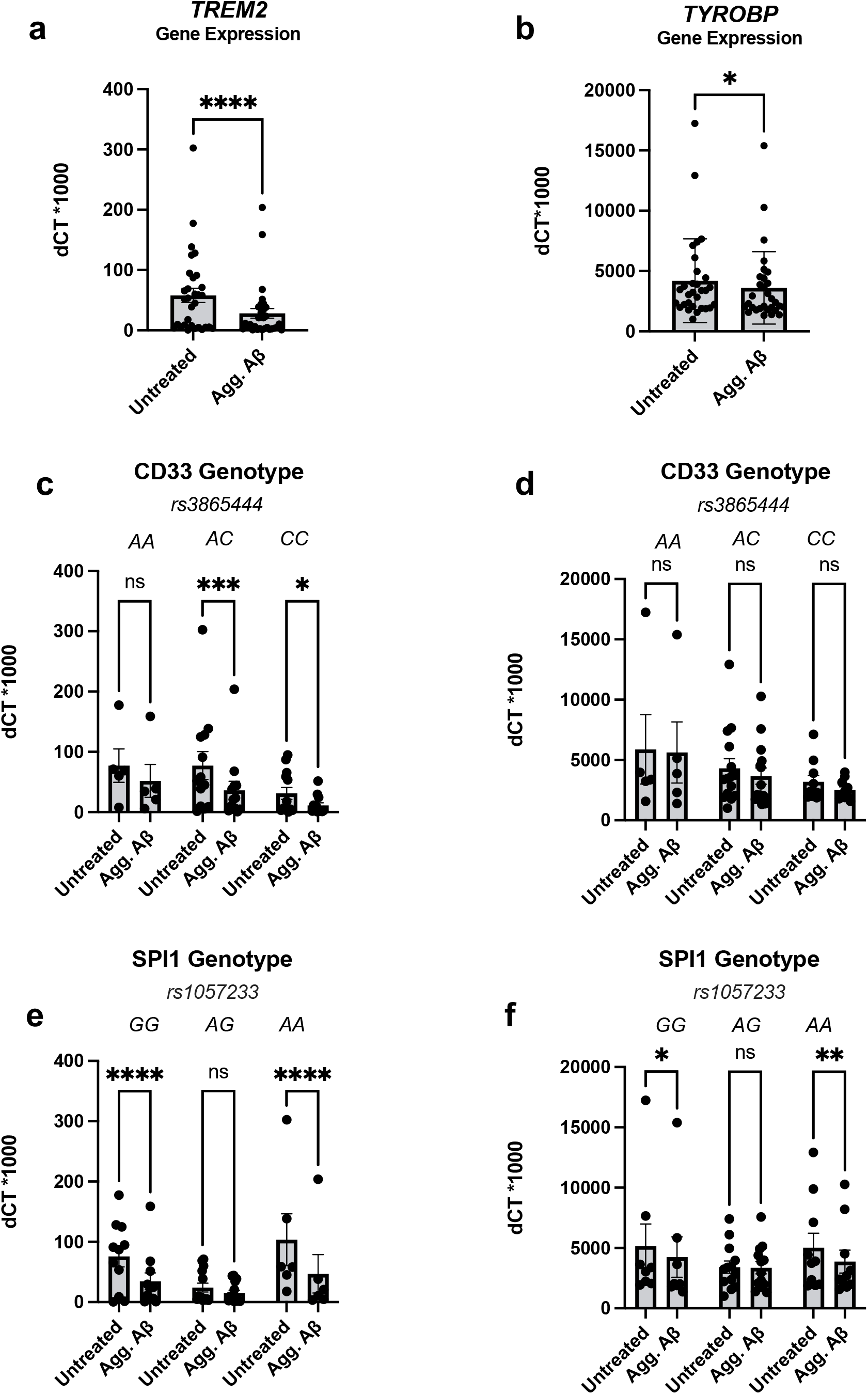
Aggregated Aβ1-42 Treatment Influences *TREM2* and *TYROBP* Transcription in a Genotype-Independent Manner. **a-b:** Human monocyte RNA expression of *TREM2***(a)** and *TYROBP***(b)** following exposure to aggregated amyloid beta peptide 1-42 (Agg. Aβ) for 24 hours across all genotypes. **c-d:** RNA expression of *TREM2***(c)** and *TYROBP***(d)** as in **a-b**, stratified by *CD33 rs3865444* genotype. *CD33 rs3865444*^*C*^ is associated with increased AD risk. **e-f:** RNA expression of *TREM2***(e)** and *TYROBP***(f)** as in **a-b**, stratified by *SPI1 rs1057233* genotype. *SPI1 rs1057233*^*A*^ is associated with increased AD risk. Gene expression results were measured using quantitative real-time PCR and were normalized to *GAPDH* expression. Results are depicted as the delta Ct value between the gene of interest and GAPDH times 1000. **a-f**: Data is presented as the mean ± SEM, and each dot represents an individual donor. **a-b**: paired Student’s t-test, **c-f**: two-way ANOVA with Sidak’s multiple comparison test *p<0.05, **p<0.01.

**Figure 4.**
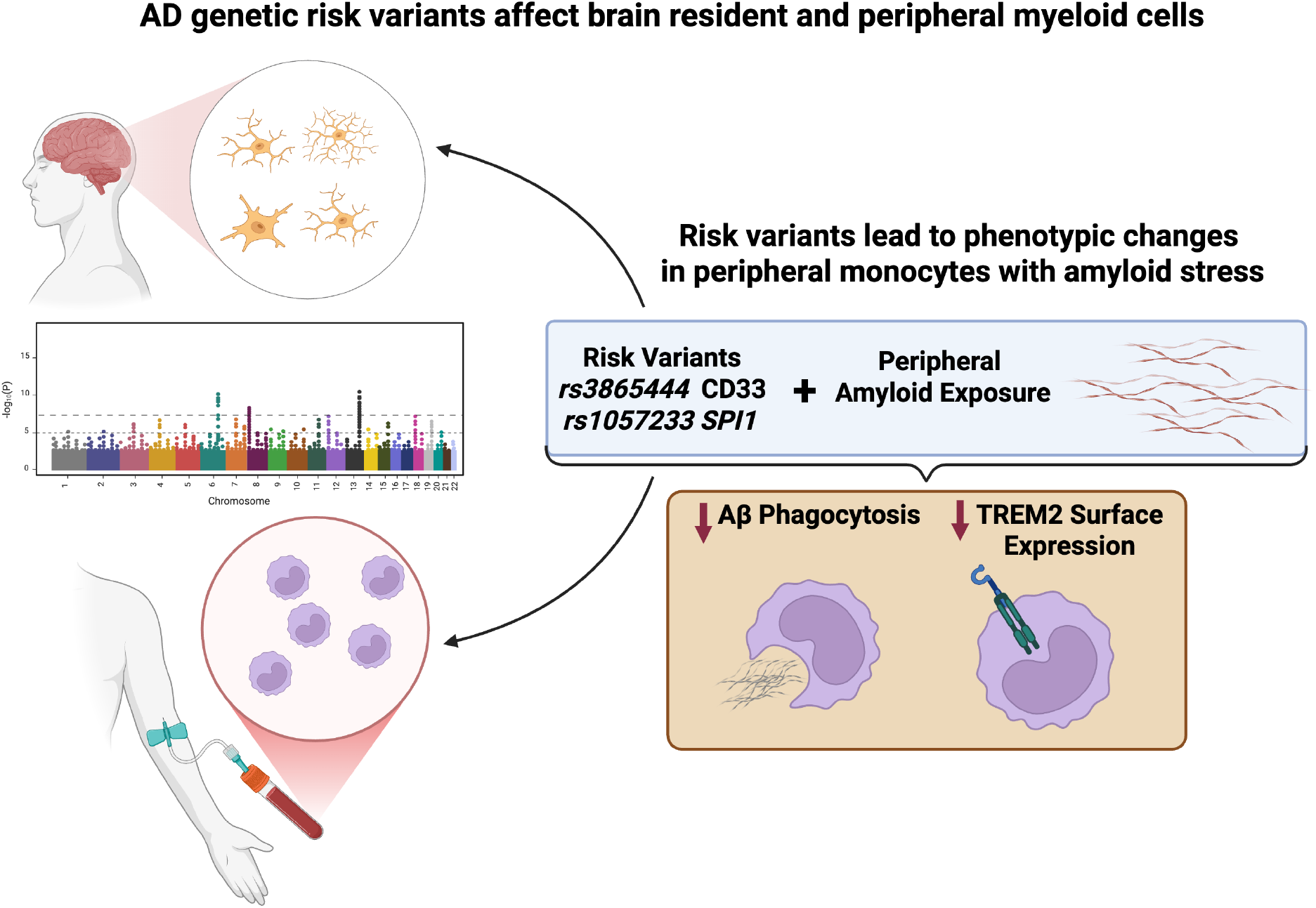
Working Model. Genetic risk variants enriched in myeloid cells impact the phenotype of microglia as well as myeloid cells in the periphery, including monocytes. Exposure to amyloid beta (Aβ) in peripheral monocytes carrying *CD33* and *SPI1* AD risk variants results in impaired phagocytosis of Aβ and reduced TREM2 surface expression.

## Discussion

In this work, we have demonstrated that aggregated Aβ1-42 treatment exposes functional impairment associated with AD risk factors in human monocytes. We found that in monocytes carrying homozygous *CD33* or *SPI1* AD risk variants, aggregated Aβ1-42 revealed deficits in phagocytosis and reduced TREM2 surface expression. Importantly, PU.1 is a transcription factor that regulates the expression of several AD-associated myeloid genes. Thus, we further investigated this pathway at the RNA level and found that aggregated Aβ1-42 reduced gene expression of *TREM2* and *TYROBP*, implicating altered regulation of this axis in the context of Aβ exposure. These findings implicate a functional interaction between TREM2, CD33, and PU.1 in monocytes, and suggest that modulating signaling downstream of TREM2 may provide a mechanism to alter functional deficits associated with these AD risk variants.

Our findings demonstrate that aggregated Aβ1-42 exposure in the context of *CD33* and *SPI1* AD risk variants unveils functional impairment in phagocytosis. Importantly, *CD33 rs3865444*^*C*^ is associated with impaired phagocytosis^7^. Our study was limited by a small sample of individuals carrying the *CD33 rs3865444*^*AA*^ genotype, limiting our ability to replicate this finding in untreated monocytes. However, the present work clearly demonstrates a dramatic superimposed deficit in phagocytosis in *CD33 rs3865444*^*CC*^ monocytes following Aβ1-42 treatment, highlighting an important gene-by-environment interaction in the context of human monocytes. *SPI1* has also been found to regulate phagocytosis, surprisingly such that knockout of *SPI1* in BV2 cells resulted in decreased phagocytosis^14^. Our findings demonstrate that the *SPI1 rs1057233*^*AA*^ risk genotype, which results in increased PU.1 expression^11^, results in decreased phagocytosis in the context of aggregated Aβ1-42 exposure. These findings highlight the importance of reconciling genotype and cell type specific effects of *SPI1 rs1057233* with knockout studies, to better understand the functional implications of this risk variant in AD.

We observed different effects of aggregated Aβ1-42 treatment on TREM2 regulation at the RNA and the surface protein level, such that while *TREM2* RNA expression was reduced in a genotype-independent manner, TREM2 surface expression was only reduced in monocytes carrying homozygous *CD33* or *SPI1* risk variants. The discordance between the mRNA TREM2 levels and surface protein expression suggests that those with homozygous risk variants alone are unable to maintain stable TREM2 surface levels in the face of reduced *TREM2* gene expression following aggregated Aβ1-42 exposure, which may have functional implications such as the impaired phagocytosis we observed. These findings also suggest that post-transcriptional regulation of TREM2 may differ in the risk variants following aggregated Aβ1-42 exposure. Importantly, TREM2 does not only exist as a surface protein but also in soluble forms, either secreted or cleaved, and can be sequestered intracellularly. The risk variants may have altered regulation between these different forms of TREM2. Future work may investigate differences in secreted or cleaved TREM2 across genotypes to better understand the consequences of *CD33* and *SPI1* genotypes on the regulation of this protein in the context of Aβ. Importantly, while PU.1 is known to regulate *TREM2* and *TYROBP* gene expression, we did not observe differences in gene expression in untreated monocytes across *SPI1 rs1057233* genotypes. The body of literature surrounding PU.1 regulation of these genes is based on knockout studies, and further work specifically investigating the impact of *SPI1 rs1057233* genotype on the regulation of these genes in all myeloid cells is needed.

Interestingly, the *CD33 rs3865444*^*C*^ risk variant is associated with increased surface TREM2 expression, an effect which we also observed in our data^8^. This *CD33* risk variant is also associated with increased CD33 surface expression, with 7-fold increased expression reported in the *rs3865444*^*CC*^ genotype compared to *rs3865444*^*AA 7*^. As CD33 is an inhibitory receptor, it is possible that increased expression of the activating TREM2 receptor in the *rs3865444*^*CC*^ risk genotype may be a compensatory mechanism to balance myeloid cell activation. The reduction of surface TREM2 levels we observed with *CD33* homozygous risk following Aβ1-42 treatment importantly occurs in the context of this increased CD33 surface expression, likely resulting in a skewed balance of myeloid cell activating receptors compared to the *rs3865444*^*AA*^ protective genotype. Alternatively, *CD33 rs3865444*^*CC*^ may limit cleavage of TREM2 into its soluble form, which has been suggested to be protective^20^. While we did not investigate the AD-associated *TREM2*^*R47H*^ variant because of its low frequency in the population, this mutation also reduces both TREM2 function and SYK activation, which aligns with the functional outcomes detailed in our findings. As the direction of effect of several AD genetic risk factors results in reduced myeloid activating tone^1^, these findings suggest that aggregated Aβ1-42 may interact with genetic background to exacerbate reduced myeloid efficacy associated with disease variants, as we observed here.

Activating molecules downstream in the TREM2 signaling pathway may reduce the phagocytosis impairment we observed associated with risk variants. TREM2, TREM1, as well as other innate immune receptors including CLEC7A, activate shared downstream signaling molecules, including SYK. Indeed, other studies have found that activating SYK through CLEC7A in mice expressing the AD-associated human *TREM2*^*R47H*^ rescued microglial activation^21^. Future work is needed to evaluate whether SYK activation is sufficient for this effect. SYK activation may be evaluated across genotypes by measuring phospho-SYK to better understand how this signaling pathway is impacted following Aβ treatment. SYK inhibitors and direct SYK activators may also be employed to elucidate the role of this signaling molecule in modulating myeloid cell phenotype in the context of Aβ.

Our work raises questions as to the role of monocyte dysfunction in AD. Monocytes have been implicated in AD through several other lines of evidence in addition to the genetic studies which suggest myeloid involvement in disease^1^. Monocytes in the CSF have been found to be dysregulated in aging^22^. Furthermore, peripheral monocytes from individuals with AD were found to have impaired Aβ phagocytosis, which aligns with the phenotype we found associated with *CD33* and *SPI1* risk variants here^23,24^. Importantly, amyloid clearance in the periphery has been linked to amyloid clearance in the brain, such that elevated amyloid in the periphery reduces clearance in the brain^4^. In addition to their role in the periphery, patrolling monocytes have been found to infiltrate the brain and differentiate into activated macrophages^25^. Furthermore, monocytes in a mouse model of AD were found to traffic to the brain to eliminate amyloid aggregates from amyloid-laden brain vasculature and then traffic back to the periphery^3^. In this model, the removal of monocytes resulted in increased amyloid burden in the cortex and hippocampus^3^. Together, this evidence suggests that monocytes may not only impact CNS amyloid dynamics both locally and from the periphery, but that their phagocytic function and phenotype is also altered in AD. Recent work has also highlighted the importance of monocytes in rescuing exhausted microglia in AD by infiltrating the CNS and acting as a new local brain myeloid population^5^.

New immunotherapies targeting TREM2 on microglia in the CNS with activating antibodies are currently being developed^15,26^. However, adverse outcomes have been reported with both anti-amyloid and anti-TREM2 antibody therapies in the CNS, including amyloid-related imaging abnormalities^27,28^. Our findings bring into question whether modulating peripheral myeloid cell activation alone may provide therapeutic benefit in AD and contribute to a foundation for future work to investigate novel therapeutic approaches that modulate the peripheral immune system.

## Methods

### Human Blood Samples

The New York Blood Center (NYBC) provided IRB-compliant access to blood samples of de-identified individuals. NYBC donors included in this study range in age from 22 to 73 years of age and include 13 females and 17 males, as detailed in **Supplemental Table 1**.

### Genotyping

Genomic DNA (gDNA) was purified from peripheral blood mononuclear cells (PBMCs) using the Gentra Puregene Blood Kit (Qiagen). DNA quantification was performed using the NanoDrop ND-1000 spectrophotometer (NanoDrop Technologies). The genotypes for *CD33 rs3865444* and *SPI1 rs1057233* were determined using Taqman SNP Genotyping Assays (Thermo Fisher Scientific Cat #4351379) on the QuantStudio 3 Real-Time PCR System. Approximately 25 ng of gDNA was used in each reaction.

The *CD33 rs3865444* context sequence used for genotyping was: CTATATCCTGCTGGACTAAACACCC[C/A]ATGGATCTAGGTGAGGCTGCGACTC

The *SPI1 rs1057233* context sequence used for genotyping was: AACTTTACTTGTTTTTTGGGAGGAG[A/G]TTAATGGGTGGGAGGGGTGAGAGGG

### Collection of Human Peripheral PBMCs and Monocyte Isolation

PBMCs were separated by Lymphoprep gradient centrifugation (StemCell Technologies). PBMCs were frozen at a concentration of 2 × 10^7^ cells ml^−1^ in 10% DMSO (Sigma-Aldrich)/90% fetal bovine serum (vol/vol, Corning). After thawing, PBMCs were washed in 10 ml of phosphate-buffered saline. Monocytes were positively selected from whole PBMCs using anti-CD14+ microbeads (Miltenyi Biotec) and plated at 2 × 10^5^ cells per well in 96-well plates.

### Amyloid Aggregation

Aβ1-42 (rPeptide) was reconstituted in 1% of NH_4_OH in MilliQ water for 10 min at RT. Then, the solution was incubated at 37°C for 1 week with rotation. Cells at a concentration of 2 × 10^5^ cells per well were incubated with 250 nM aggregated Aβ1-42 in RPMI medium for 24 hours at 37°C.

### Amyloid Phagocytosis Assay

Amyloid-beta 1-42 phagocytosis assays were performed according to a previously published protocol^7^. Briefly, monocytes were incubated with 15 ng ml^−1^ HiLyte Fluor 647–labeled Aβ1-42 (AnaSpec) in 96-well polypropylene plates in RPMI media plus 5% fetal calf serum (vol/vol) for 1 h at 37 °C or 4 °C. As a negative control, cells were preincubated with 10µM Cytochalasin D (FisherSci.) for 30 minutes before being treated with 647–labeled Aβ1-42. Cells were then washed three times with PBS with 1% fetal calf serum and labeled with Fixable LIVE/DEAD violet cell stain (Thermo Fisher Scientific) for 30 minutes and fixed in 4% paraformaldehyde (vol/vol) for 30 min. Flow cytometry data were collected on a NovoCyte Quanteon cell analyzer and compensated and analyzed using the associated NovoExpress software.

### TREM2 Surface Staining

Cells were stained on ice with TREM2-APC (R&D Systems, 1:20) for 30 min protected from light, according to the manufacturer. Isotype controls (R&D Systems) were similarly stained. Cells were then washed three times with PBS with 1% fetal calf serum and labeled with Fixable LIVE/DEAD violet cell stain (Thermo Fisher Scientific) for 30 minutes and fixed in 4% paraformaldehyde (vol/vol) for 30 min. Flow cytometry data were collected on a NovoCyte Quanteon cell analyzer and compensated and analyzed using the associated NovoExpress software.

### RNA Expression of *TREM2* and *TYROBP*

Monocyte populations were exposed to aggregated Aβ1-42 for 24 hours and were subsequently lysed for RNA isolation, which was performed using the RNAeasy isolation kit (Qiagen, cat#74134). cDNA was made from isolated RNA using the High Capacity cDNA Reverse Transcription Kit (ThermoFisher), according to the manufacturer’s protocol. Quantitative real-time PCR (qRT-PCR) was performed using Taqman probes for each of the assayed genes and Taqman Fast Advanced Mastermix (ThermoFisher). qRT-PCR was run on a QuantStudio3 (ThermoFisher) and analyzed using the associated Design and Analysis Software. A *GAPDH* TaqMan probe (ThermoFisher) was used as a housekeeping gene. Relative gene expression was calculated as the delta CT values between the gene of interest and *GAPDH* expression, times 1000.

### Batch correction

Batches were normalized in GraphPad Prism using the normalize function. Briefly, the function averaged and normalized the means of each condition. 0% was defined as the smallest mean in each dataset, and 100% was defined as the largest mean in each dataset, and the results were presented as fractions.

### Statistical Analysis

Statistical analyses were performed using GraphPad Prism (GraphPad Software). Student’s *t*-test was performed for comparisons between individual groups. For group-wise comparison, two-way-ANOVA was used with Sidak’s multiple comparison test. A *p*-value less than 0.05 was considered significant.

## Acknowledgements

This work was supported by the US National Institutes of Health grants RF1AG058852, F30AG074618, and R21AG073882, as well as Alzheimer’s Association grant DVT-14-321148. The content is solely the responsibility of the authors and does not necessarily represent the official views of the National Institutes of Health. EMB is a current Ludwig Scholar in the Carol and Gene Ludwig Center for Research in Neurodegeneration. Flow cytometry was performed in the Columbia Stem Cell Initiative Flow Cytometry core facility at Columbia University Irving Medical Center under the leadership of Michael Kissner.

