## Extended Data for "Alzheimer’s Disease Risk Variants Interact with Amyloid-beta to Modulate Monocyte Function"

a

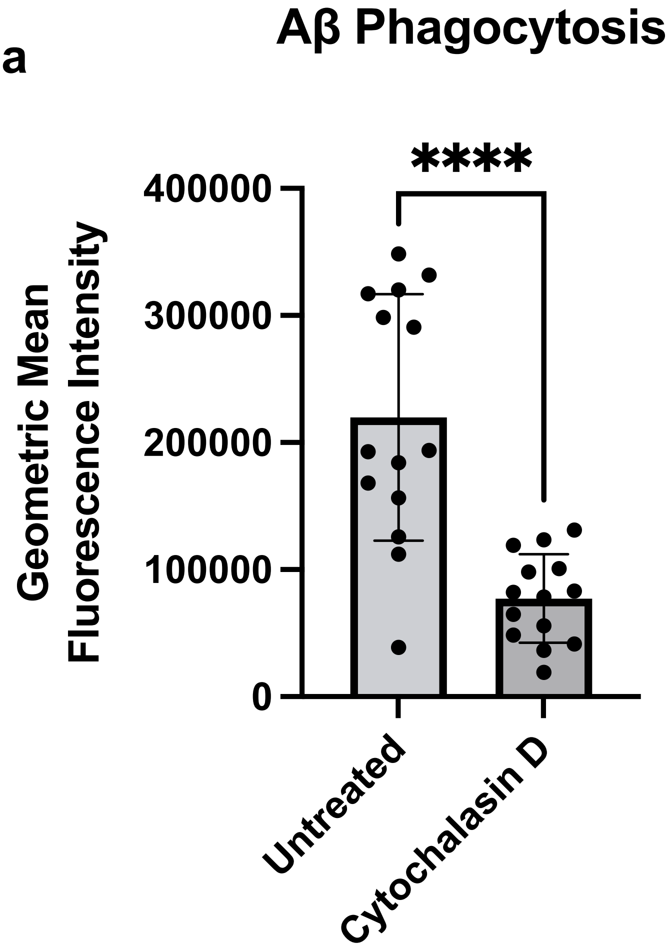
Aβ Phagocytosis
****
Geometric Mean
Fluorescence Intensity
400000
300000
200000
100000
0
Untreated
Cytochalasin D

b

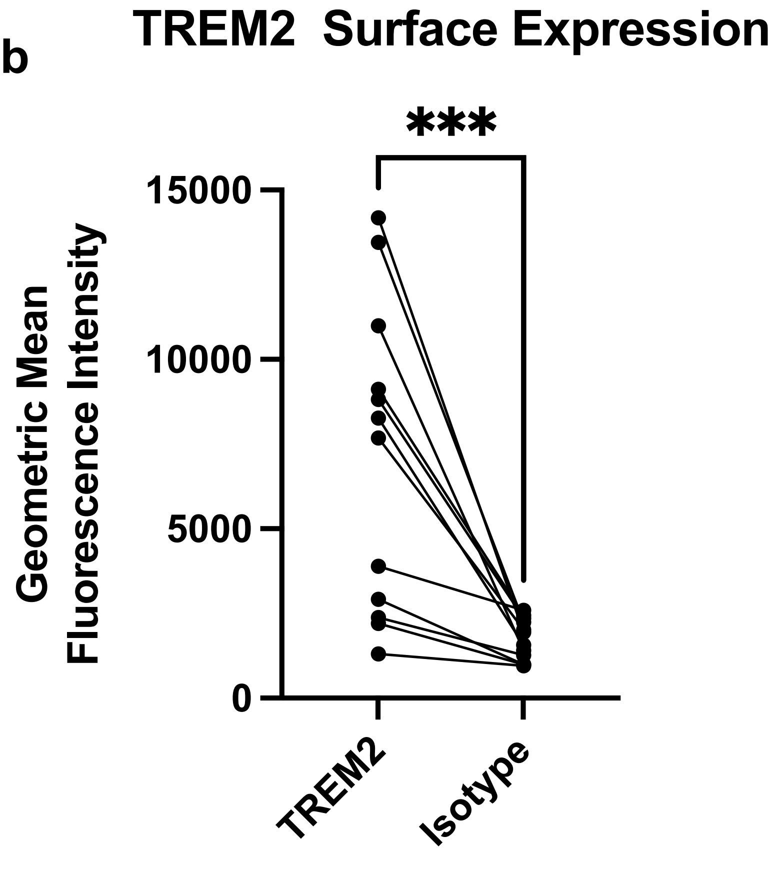
TREM2 Surface Expression
***
Geometric Mean
Fluorescence Intensity
15000
10000
5000
0
TREM2
Isotype

**Extended Data Figure 1. Phagocytosis and Surface Staining Controls a.** Monocyte phagocytosis of fluorescently labeled amyloid beta peptide 1-42 following incubation with cytochalasin D for 30 minutes. **b** Monocyte surface expression of TREM2 or isotype control. All data were collected with flow cytometry and represented as geometric mean fluorescence intensity. Each dot represents an individual donor. For **a** data is presented as mean $\pm$ SEM, **b** paired samples are connected by a line. Paired Student’s t-test, ***$p<0.001$, ****$p<0.0001$.

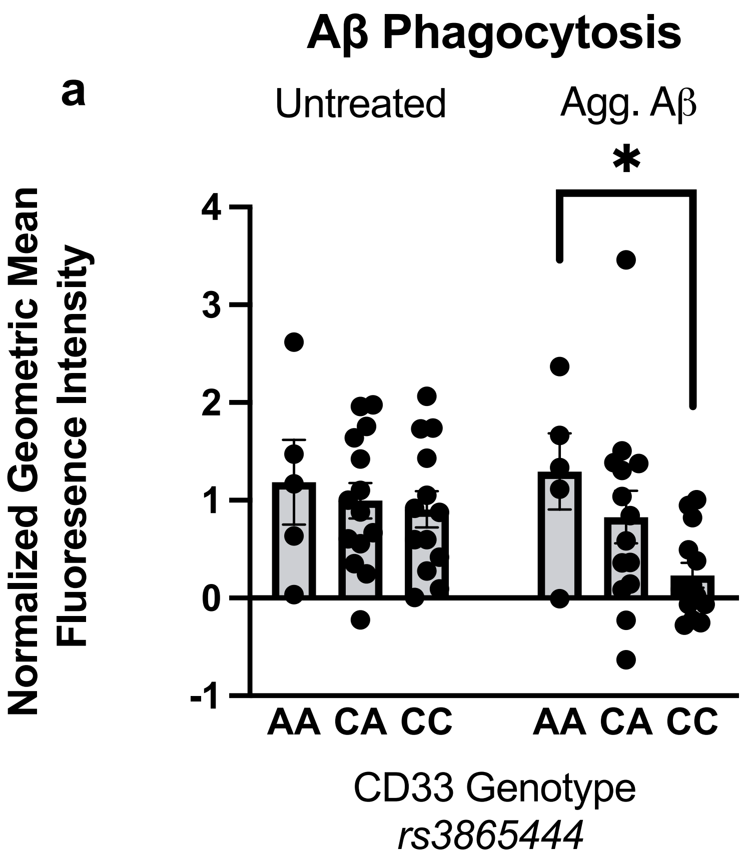
Aβ Phagocytosis
a
Untreated
Agg. Aβ
*
Normalized Geometric Mean Fluoresence Intensity
4
3
2
1
0
-1
AA CA CC AA CA CC
CD33 Genotype
rs3865444

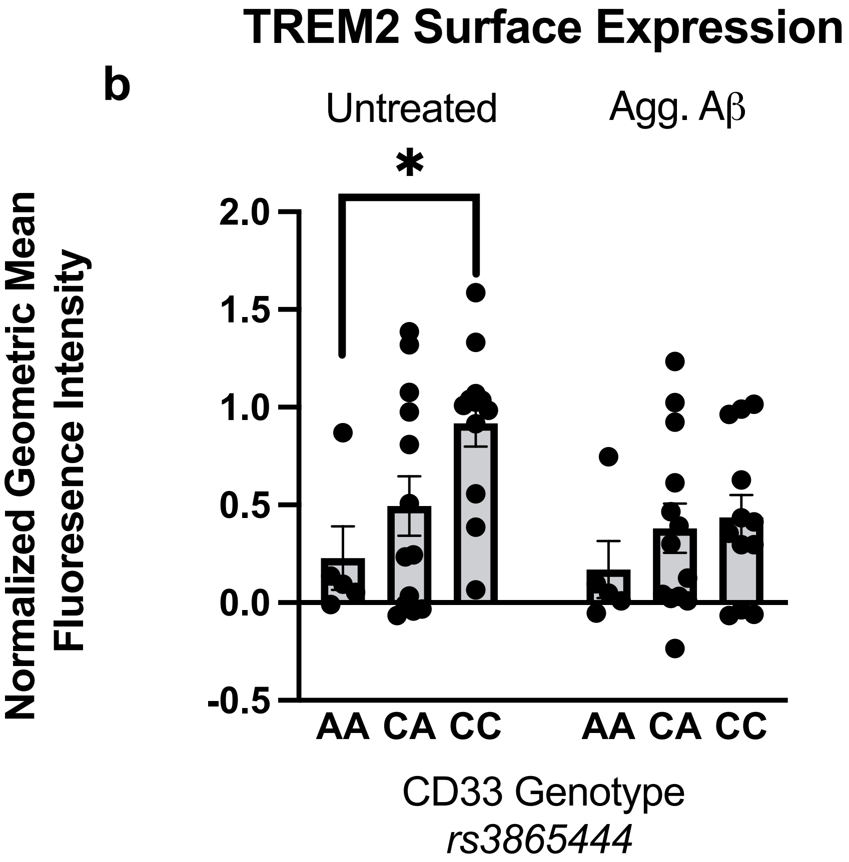
TREM2 Surface Expression
b
Untreated
Agg. Aβ
*
Normalized Geometric Mean Fluoresence Intensity
2.0
1.5
1.0
0.5
0.0
-0.5
AA CA CC AA CA CC
CD33 Genotype
rs3865444

**Extended Data Figure 2. Aβ Phagocytosis and TREM2 Surface Expression by *CD33 rs3865444* Genotype.** **a**. Monocyte phagocytosis of fluorescently labeled amyloid beta peptide 1-42 (Aβ1-42) following exposure to aggregated Aβ1-42 (Agg. Aβ) for 24 hours in *CD33 rs3865444* genotype groups. **b.** Monocyte surface expression of TREM2 following exposure to aggregated Aβ1-42 (Agg. Aβ) for 24 hours in *CD33 rs3865444* genotype groups. All data were measured with flow cytometry, represented as geometric mean fluorescence intensity and normalized for batch correction using the GraphPad Prism normalize function. Data is presented as the mean ± SEM and each dot represents an individual donor. Two-way ANOVA with Sidak's multiple comparison test *$p<0.05$.

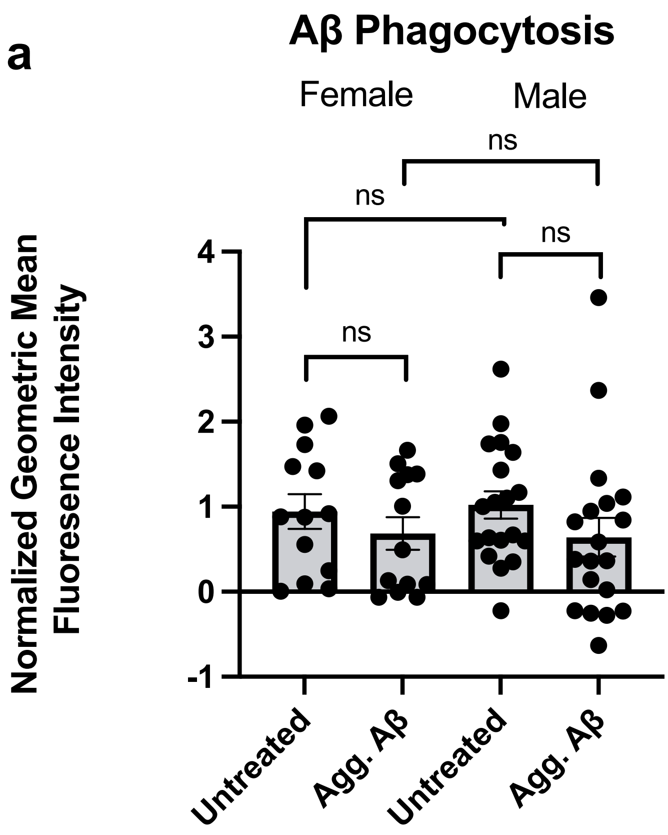
a
Aβ Phagocytosis
Female
Male
ns
ns
ns
ns
Normalized Geometric Mean
Fluoresence Intensity
4
3
2
1
0
-1
Untreated
Agg. Aβ
Untreated
Agg. Aβ

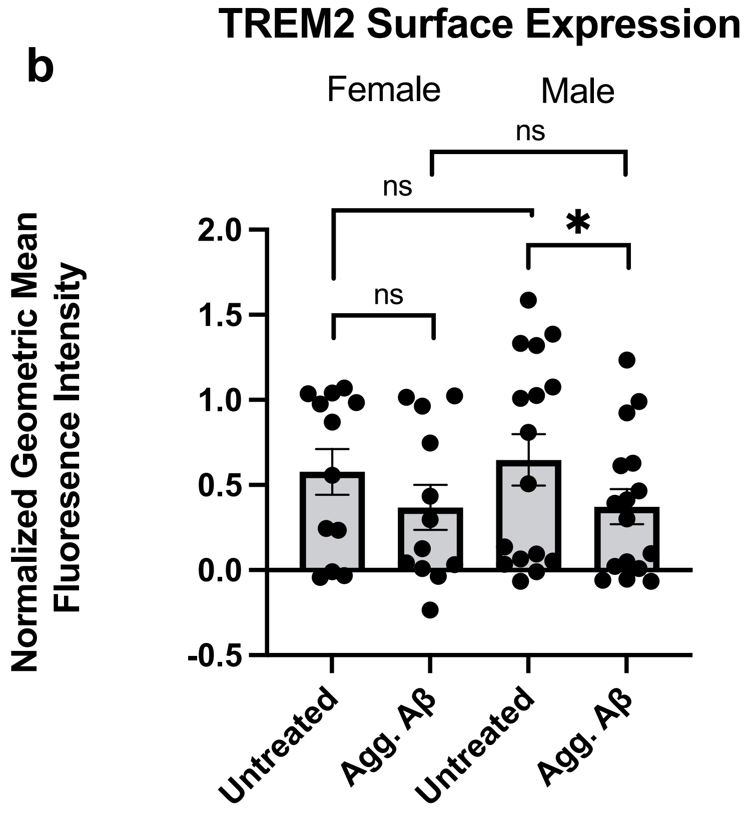
b
TREM2 Surface Expression
Female
Male
ns
ns
*
ns
Normalized Geometric Mean
Fluoresence Intensity
2.0
1.5
1.0
0.5
0.0
-0.5
Untreated
Agg. Aβ
Untreated
Agg. Aβ

**Extended Data Figure 3. Aβ Phagocytosis and TREM2 Surface Expression Stratified by Sex. a**. Monocyte phagocytosis of fluorescently labeled amyloid beta peptide 1-42 (Aβ1-42) following exposure to aggregated Aβ1-42 (Agg. Aβ) for 24 hours in males and females. **b.** Monocyte surface expression of TREM2 following exposure to aggregated Aβ1-42 (Agg. Aβ) for 24 hours in males and females. All data were measured with flow cytometry, represented as geometric mean fluorescence intensity and normalized for batch correction using the GraphPad Prism normalize function. Data is presented as the mean $\pm$ SEM and each dot represents an individual donor. Two-way ANOVA with Sidak's multiple comparison test *$p<0.05$.

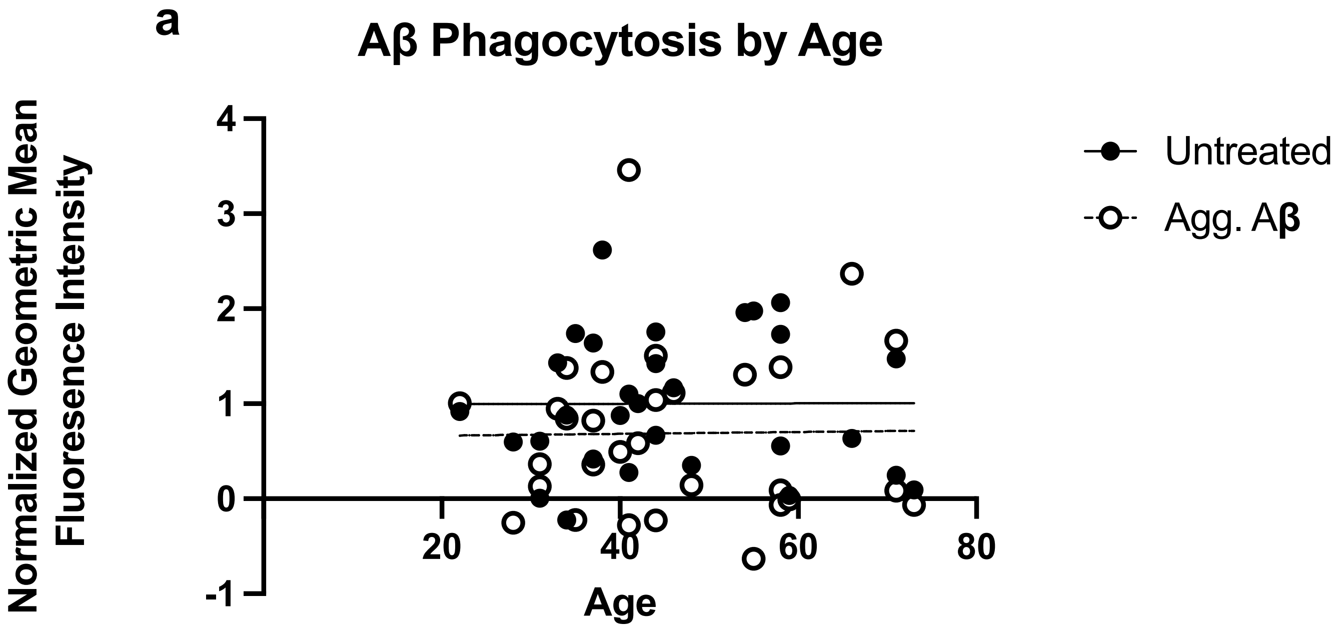
a
Aβ Phagocytosis by Age
Normalized Geometric Mean Fluoresence Intensity
4
3
2
1
0
-1
20
40
60
80
Age
Untreated
Agg. Aβ

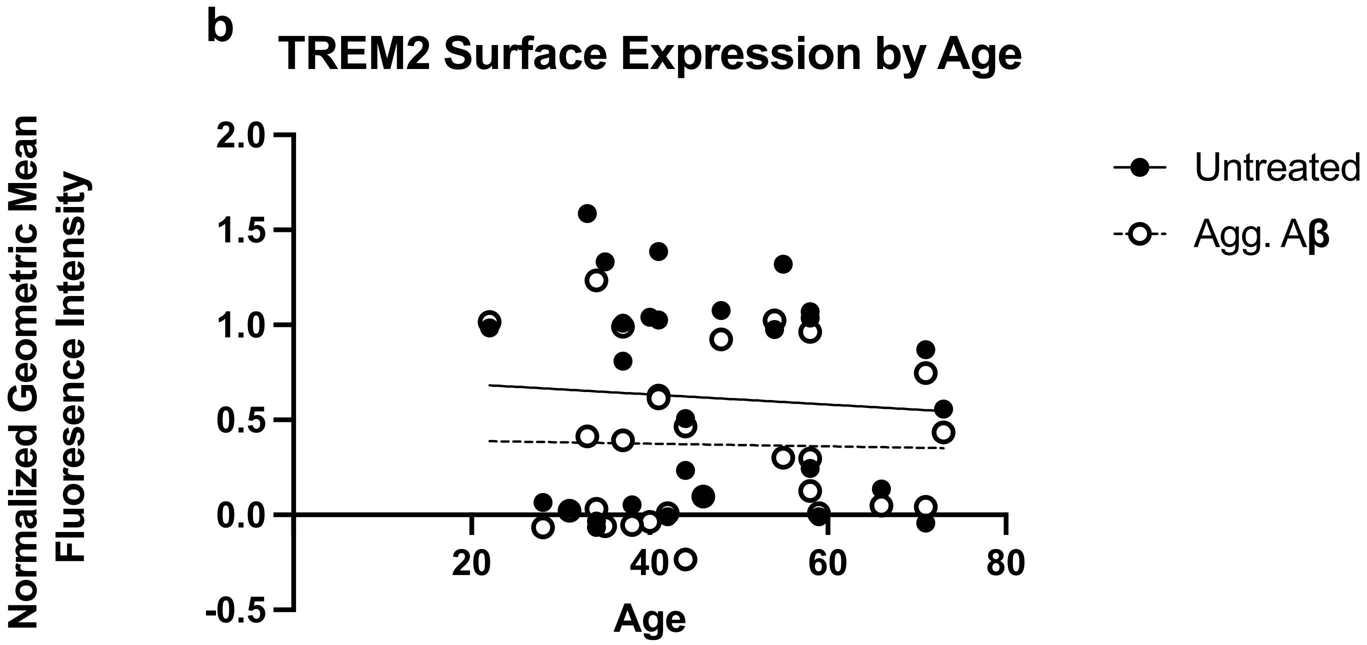
b
TREM2 Surface Expression by Age
Normalized Geometric Mean Fluoresence Intensity
2.0
1.5
1.0
0.5
0.0
-0.5
20
40
60
80
Age
Untreated
Agg. Aβ

**Extended Data Figure 4. Aβ Phagocytosis and TREM2 Surface Expression by Age. a**. Monocyte phagocytosis of fluorescently labeled amyloid beta peptide 1-42 (Aβ1-42) following exposure to aggregated Aβ1-42 (Agg. Aβ) for 24 hours versus age. **b.** Monocyte surface expression of TREM2 following exposure to aggregated Aβ1-42 (Agg. Aβ) for 24 hours versus age. All data were measured with flow cytometry, represented as geometric mean fluorescence intensity and normalized for batch correction using the GraphPad Prism normalize function. Simple linear regression.

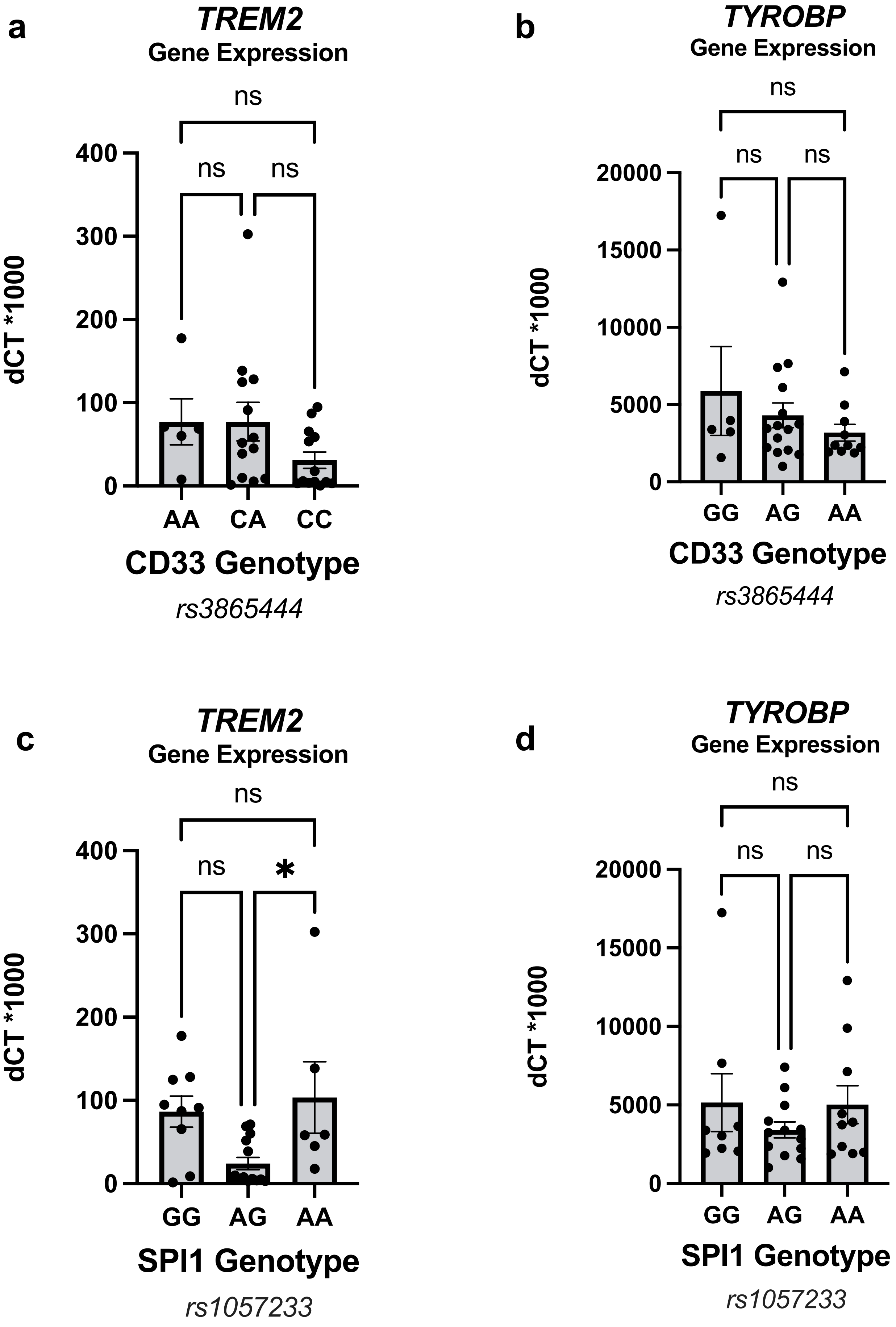

a
TREM2
Gene Expression
ns
ns
ns
400
300
200
100
0
dCT *1000
AA
CA
CC
CD33 Genotype
rs3865444
b
TYROBP
Gene Expression
ns
ns
ns
20000
15000
10000
5000
0
dCT *1000
GG
AG
AA
CD33 Genotype
rs3865444
c
TREM2
Gene Expression
ns
ns
*
400
300
200
100
0
dCT *1000
GG
AG
AA
SPI1 Genotype
rs1057233
d
TYROBP
Gene Expression
ns
ns
ns
20000
15000
10000
5000
0
dCT *1000
GG
AG
AA
SPI1 Genotype
rs1057233

**Extended Data Figure 5. Baseline Gene Expression of TREM2 and TYROBP by *CD33 rs3865444* and *SPI1 rs1057233* Genotypes.** RNA expression of *TREM 2***(a,c)** and *TYROBP* **(b,d)** in untreated human monocytes. **a**. *TREM2* RNA expression stratified by *CD33 rs3865444* genotype. **b**. *TYROBP* RNA expression stratified by *CD33 rs3865444* genotype. **(c)** *TREM2* RNA expression stratified by *SPI1 rs1057233* genotype. **(d)** *TYROBP* RNA expression stratified by *SPI1 rs1057233* genotype. Gene expression results were measured using quantitative real-time PCR and were normalized to *GAPDH* expression. Results are depicted as the delta CT value between the gene of interest and GAPDH times 1000. Data is presented as the mean ± SEM, and each dot represents an individual donor.
One-way ANOVA with Sidak's multiple comparison test *$p<0.05$.
